# SARS-CoV-2 Entry Protein TMPRSS2 and Its Homologue, TMPRSS4 Adopts Structural Fold Similar to Blood Coagulation and Complement Pathway Related Proteins

**DOI:** 10.1101/2021.04.26.441280

**Authors:** Vijaykumar Yogesh Muley, Amit Singh, Karl Gruber, Alfredo Varela-Echavarría

## Abstract

The severe acute respiratory syndrome coronavirus 2 (SARS-CoV-2) utilizes TMPRSS2 receptor to enter target human cells and subsequently causes coronavirus disease 19 (COVID-19). TMPRSS2 belongs to the type II serine proteases of subfamily TMPRSS, which is characterized by the presence of the serine-protease domain. TMPRSS4 is another TMPRSS member, which has a domain architecture similar to TMPRSS2. TMPRSS2 and TMPRSS4 have been shown to be involved in SARS-CoV-2 infection. However, their normal physiological roles have not been explored in detail. In this study, we analyzed the amino acid sequences and predicted 3D structures of TMPRSS2 and TMPRSS4 to understand their functional aspects at the protein domain level. Our results suggest that these proteins are likely to have common functions based on their conserved domain organization. Furthermore, we show that the predicted 3D structure of their serine protease domain has significant similarity to that of plasminogen which dissolves blood clot, and of other blood coagulation related proteins. Additionally, molecular docking analyses of inhibitors of four blood coagulation and anticoagulation factors show the same high specificity to TMPRSS2 and TMPRSS4 3D structures. Hence, our observations are consistent with the blood coagulopathy observed in COVID-19 patients and their predicted functions based on the sequence and structural analyses offer avenues to understand better and explore therapeutic approaches for this disease.

## 1. Introduction

Proteolysis is mediated by a special class of proteins called proteases or peptidases that hydrolyze peptide bonds of their substrate proteins (López-Otín and Overall, 2002). They act as a surveillance system that monitors the turnover of cellular proteins. Hence, they modulate a plethora of cellular processes including cell growth, survival, and death, as well as phagocytosis, signaling pathways and membrane re-modelling (Muley et al., 2019; Puente et al., 2005). In *Escherichia coli*, 36% (26% with stringent criteria) of proteases belong to the serine protease family (Clausen et al., 2002) and this distribution is estimated to be similar for many organisms. More than two percent of human genes encode proteases (Puente et al., 2005), and 20 of them are classified as the type II transmembrane serine proteases (TTSP). TTSPs have conserved domain organization, which consists of a single-pass transmembrane domain located near the amino-terminal end of the protein spanning through the cytosol and a large extracellular portion at the carboxy-terminus containing the serine protease domain of the chymotrypsin fold (Clausen et al., 2002; Szabo and Bugge, 2008). This fold is characterized by the Ser-His-Asp catalytic triad, which is involved in endopeptidase activity. These enzymes are widely distributed in prokaryotic and eukaryotic genomes (Clausen et al., 2002; Muley et al., 2019; Puente et al., 2005). Interestingly, the first TTSP member was identified over a century ago by Pavlov due to its essential role in food digestion (Szabo and Bugge, 2008), and it was cloned in 1994 leading to its characterization as a plasma membrane-anchored protein (Kitamoto et al., 1994).

The transmembrane protease, serine 2 (TMPRSS2) and 4 (TMPRSS4) are members of the TTSP family and belong to the hepsin/transmembrane protease/serine (TMPRSS) subfamily of TTSP (Szabo and Bugge, 2008). TMPRSS2 facilitates SARS-CoV-1 and SARS-CoV-2 entry in human cells and plays a critical role in Coronavirus disease 19 (Covid19) (Hoffmann et al., 2020; Hu et al., 2020; Matsuyama et al., 2010). TMPRSS4 was previously characterized as TMPRSS3 (Wallrapp et al., 2000), which along with TMPRSS2 promotes SARS CoV-2 infection in human enterocytes (Zang et al., 2020). Its overexpression has been observed in dozens of cancers and it contributes to tumorigenesis and metastasis (Aberasturi and Calvo, 2015; Lee et al., 2016; Villalba et al., 2019). Interestingly, TMPRSS2 and TMPRSS4 have been also shown to act as host cell entry receptors for Influenza virus (Bertram et al., 2010) and TMPRSS2 was further shown to be involved in replication of H7N9 and Influenza viruses *in vivo* (Sakai et al., 2014). However, their functions are not clearly understood in normal conditions or in viral diseases.

In this study, we analyzed the amino acid sequences and predicted 3D structures of TMPRSS2 and TMPRSS4 to understand their functional aspects at the protein domain level. Our results suggest that these proteins are likely to have common functions based on their conserved domain organization. Furthermore, we show that the predicted 3D structure of their serine protease domain has significant similarity to that of plasminogen, and of other blood coagulation related proteins. Additionally, molecular docking analyses of inhibitors of four blood coagulation and anticoagulation factors show the same high specificity to TMPRSS2 and TMPRSS4 3D structures. Hence, our observations are consistent with the blood coagulopathy observed in Covid19 patients and their predicted functions based on the sequence and structural analyses offer avenes to understand better and explore therapeutic approaches for this disease.

## 2. Material and methods

### 2.1. Sequence analysis

Protein sequences of TMPRSS2 and TMPRSS4 from humans and their mouse orthologs were obtained from the UniProt database (Bateman et al., 2017). Protein domains were identified using the scanProsite tool from the ProSite database (Castro et al., 2006; Sigrist et al., 2009). Further domain architecture information was obtained from the Genome3D database (Lewis et al., 2015). The TOPCONS web server was used to predict the membrane-spanning region of the proteins (Tsirigos et al., 2015). Multiple sequence alignment of human and mouse proteins was constructed using the MAFFT plugin of JalView program, and visualized using the latter (Katoh et al., 2018; Waterhouse et al., 2009). The sequences of TMPRSS2 and TMPRSS4 were used for searches against the Protein Data Bank (PDB) database using HHPred to find their structural homologs (Berman, 2000; Hildebrand et al., 2009). Phyre2 was used in intensive mode to predict their 3D structures (Kelley et al., 2015). Phyre2 modelled the TMPRSS2 structure using the PDB template structures 4O03_A, 2XRC_D, 6ESO_A, 4DUR_A, 4HZH_B, 1Z8G_A, and 3NXP_A. The same templates were also used to model the TMPRSS4 structure except the 3NXP_A. The regions composed of the scavenger receptor cysteine-rich (SRCR) and serine protease domains in TMPRSS2 and TMPRSS4 were modelled with high accuracy by Phyre2, which was also supported by HHPred results. The predicted structures belonging to this region were then uploaded to the CATH web server to obtain the structural domain hits from available crystal structures (Dawson et al., 2017). CATH results confirmed the presence of two distinct domains, a large domain with Greek-key *β*-barrel fold (Chymotrypsin domain) and a SRCR domain. Then, the 3D protein structure corresponding to this region was compared with the template structures identified by Phyre2 and top 20 structural homologs obtained from HHPred search, together containing 36 unique structures. The domain architectures of the corresponding proteins were extracted using ProSite database (Sigrist et al., 2009).

### 2.2. Protein 3D structure analysis

We computed the root mean square deviation (RMSD) between the backbone structure of the protease domain alone, the SRCR domain alone and both domains of TMPRSS2 and TMPRSS4 with the above-mentioned 36 PDB structures using the align module in PyMOL, with maximum iteration cycles of 20 and BLOSUM62 as a scoring matrix (Schrödinger, LLC, 2015). The structures of plasminogen (PDB accession, 5UGG) and prothrombin activator (a catalytic domain of prothrombinase, PDB accession, 4BXW) are available in complex with their selective inhibitors YO (trans-4-aminomethylcyclohexanecarbonyl-l-tyrosine-n-octylamide, PDB accession, 89M) and L-Glu-Gly-Arg chloromethyl ketone (PDB accession, 0GJ) respectively (Law et al., 2017; Lechtenberg et al., 2013). These structures were superimposed with TMPRSS2 and TMPRSS4 in the presence and absence of their inhibitors using the PyMOL align tool. We selected 89M and 0GJ ligands, and also the thrombin inhibitor D-phenylalanyl-N-(3-chlorobenzyl)-L-prolinamide (PDB accession, 22U) from PDB structure 2ZC9 (Baum et al., 2009) and the plasma kallikrein inhibitor, N-[(6-amino-2,4-dimethylpyridin-3-yl)methyl]-1-({4-[(1H-pyrazol-1-yl)methyl]phenyl}methyl)-1H-pyrazole-4-carboxamide (PDB accession, 75D) from 6O1S (Partridge et al., 2019) to perform their docking studies on TMPRSS2, TMPRSS4, 4BXW, 5UGG, and 2ANY structures using the Glide docking program (Friesner et al., 2004). Briefly, the LigPrep module in Maestro was employed to generate multiple conformations of the ligands followed by energy minimization (Schrödinger, 2018). The target protein structures were preprocessed to remove the bad contacts using the wizard integrated into Maestro. The OPLS force field was used to minimize the protein structure (Jorgensen and Tirado-Rives, 1988). The center of the receptor grid was placed on the center of mass (the active site triad Ser-His-Asp) of the proteins, followed by the extra precision Glide docking (Friesner et al., 2006). All structural images were rendered using PyMOL.

## 3. Results

### 3.1. A conserved extracellular domain architecture of TMPRSS2 and TMPRSS4 suggests their related functions

The human TMPRSS2 gene encodes a 492 amino acid long protein compared to the 437 amino acids encoded by TMPRSS4 (Wallrapp et al., 2000). Pairwise global sequence alignment using the Needleman-Wunsch algorithm showed amino acid similarity of 42.2% and identity of 30.3% between them (Madeira et al., 2019; Needleman and Wunsch, 1970). The single-pass transmembrane helix is present in both proteins near their N-termini (Supplementary figure 1). The approximate location of a transmembrane helix in TMPRSS2 is between 85 and 105 residues, whereas between 33 and 53 in TMPRSS4 leaving a longer N-terminal sequence of 84 amino acids in TMPRSS2. The N-terminal sequence preceding the transmembrane helix in both proteins is shorter and predicted to be cytoplasmic, while the following sequence is longer and exposed to the extracellular milieu owing to its extracellular topology (Supplementary figure 1). As shown in Figure 1A, both proteins have a conserved extracellular domain architecture containing the Low-density lipoprotein (LDL) receptor class A domain (denoted as LDLRA), the scavenger receptor cysteine rich (SRCR) and the protease domain of Peptidase S1A, chymotrypsin family or Trypsin-like serine protease superfamily according to Pfam and Superfamily database, respectively (Gough et al., 2001; Sonnhammer et al., 1997). The protease domain, hereafter referred to as serine-protease, adopts a chymotrypsin type structural fold characterized by Greek-key *β*-barrels. The results pertaining to the extracellular portion are consistent with the previous sequence analysis reports (Aberasturi and Calvo, 2015; Szabo and Bugge, 2008; Wallrapp et al., 2000). The chymotrypsin fold is a prototype structural feature of the high temperature requirement A (HtrA) protein family of Trypsin-like serine proteases, which acts as chaperones and are responsible for maintaining protein tertiary structure at high temperature (Clausen et al., 2002).

**Figure 1:**
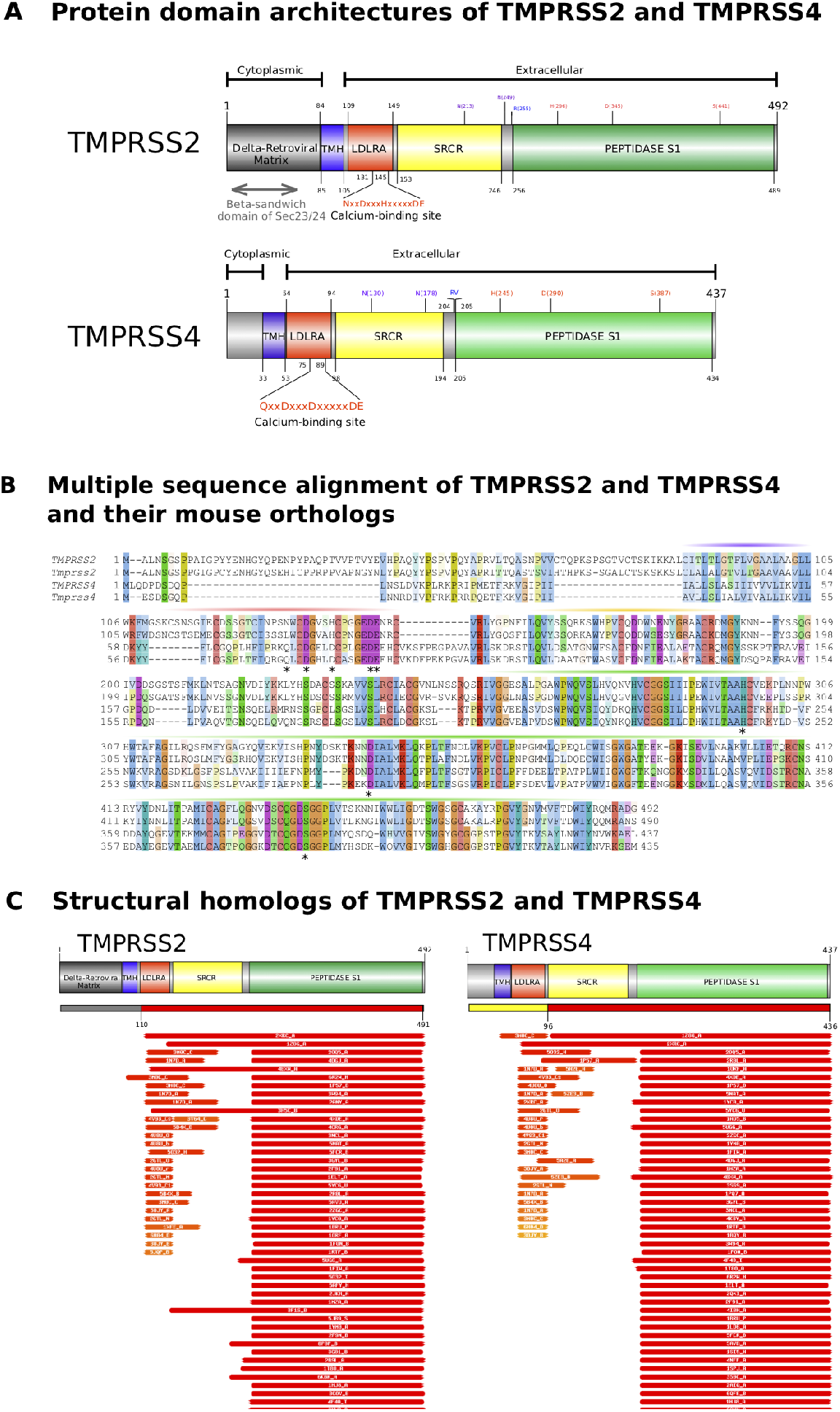
Sequence comparison of TMPRSS2 and TMPRSS4 with their mouse orthologs Tmprss2 and Tmprss4. Panel A shows domain architectures of TMPRSS2 and TMPRSS4 proteins. The following domains with known activity are shown with their amino acid positions: Low-density lipoprotein receptor class A (LDLRA_2), Scavenger receptor cysteine rich (SRCR_2), and Serine proteases, trypsin family (Trypsin). Multiple sequence alignment between human TMPRSS2 and TMPRSS4 with their mouse orthologs is shown in panel (B). The approximate location of the domains shown in (A) are indicated by bars on top of the multiple alignment in (B) with the same color code in (A). The location of the triad of Ser-His-Asp, responsible for the proteolytic activity of the Trypsin domain is conserved in all four sequences and indicated by asterisks in (B). The serine protease domain and its active sites are conserved in all proteins. The LDLRA_2 domain appears to be functional in TMPRSS2 but it is truncated in TMPRSS4, which may have the cholesterol transport activity. Panel (C) shows the structural homologs of TMPRSS2 and TMPRSS4 in the PDB database.

In contrast to conserved C-terminal regions, the N-terminal amino acid sequences preceding the transmembrane helix in both proteins differ substantially (Figure 1A). TMPRSS4 has only a stretch of 32 residues at its N-terminus, which do not show similarity to any known structure. In TMPRSS2, however, the equivalent N-terminal region of approximately 85 amino acids is predicted to be structurally homologous to the Delta-retroviral matrix superfamily (CATH superfamily code 1.10.185.10), and within this region, a stretch of amino acids spanning positions 5 to 33 show similarity to the *β*-sandwich domain of Sec23/24 superfamily (CATH superfamily code 2.60.40.1670) according to the Genome3D database annotation (Lewis et al., 2015). The *β*-sandwich domain of Sec23/24 can also be confirmed with the Superfamily database searches (SUPERFAM-ILY/SCOP database accession: 81995) (Gough et al., 2001). This domain is likely to adopt the Human T-cell Leukemia Virus Type II Matrix Protein structural fold according to CATH database annotation (Dawson et al., 2017). However, we could not detect homologous sequences for this region in viral genome restricted searches using a PSI-BLAST at NCBI, neither HHPred search predicted similar structures in the PDB database (Hildebrand et al., 2009; Johnson et al., 2008). Therefore, experimental analyses are required to address the functional aspects of both predicted domains.

### 3.2. Multiple sequence alignment of TMPRSS2 and TMPRSS4 with their mouse orthologs shows highly conserved SRCR and serine protease domains

To understand the amino acid variations in both proteins, we performed multiple sequence alignment of TMPRSS2 and TMPRSS4 with their mouse orthologs only, since our aim was to confirm whether important amino acid positions are conserved in both proteins. This analysis revealed indels in the N-terminal region of TMPRSS2 and TMPRSS4 not affecting their conserved transmembrane helix, which is followed by a conserved C-terminal sequence (Figure 1B). The LDLRA domain is located right next to the membrane helix in both proteins, which is consistent with previous studies (Aberasturi and Calvo, 2015; Szabo and Bugge, 2008). This domain contains six cysteine disulfide-bonds that bind lipoproteins such as LDLs and a highly conserved cluster of negatively charged amino acids (Bieri et al., 1995; Yamamoto et al., 1984). All six cysteines are conserved in TMPRSS2 and its mouse ortholog, and four are also conserved in TMPRSS4. One indel is adjacent to the LDLRA domain at the N-terminal end in TMPRSS4 and another at the C-terminus of TMPRSS2. The one in TMPRSS4 corresponds to one of its missing cysteine residues, and another cysteine residue is substituted by phenylalanine. Both proteins, however, have conserved calcium binding sites within this LDLRA domain. The bound calcium ion imparts structural integrity to the domain (Bieri et al., 1995), suggesting the presence of LDLRA domain activity in both proteins. This domain in both proteins is followed by a highly conserved SRCR and the serine protease domain (Figure 1B). The biochemical functions of SRCR domains have not been established with certainty but they are likely to mediate protein-protein interactions and ligand binding (Hohenester et al., 1999; Resnick et al., 1994). This domain is found in diverse secreted and membrane bound proteins including regulators of the complement cascades involved in immune response (Freeman et al., 1990). The catalytic triad of Ser-His-Asp residues responsible for its proteolytic activity is conserved in human and mouse (Figure 1B).

Overall, these results reveal that the extracellular region of TMPRSS2 and TMPRSS4, and its domain organization is highly conserved suggesting that they have related functions.

### 3.3. TMPRSS2 and TMPRSS4 show homology with complement system and blood coagulation and anticoagulation related proteins

To identify known structural homologs of both proteins, we queried their sequences in the PDB database using the HHPred webserver. The region between 110 to 491 and 96 to 436 amino acid positions in TMPRSS2 and TMPRSS4, respectively, showed significant similarity with several PDB structures (hits) (Figure 1C). We selected the top 20 significant hits for each protein for further analysis (details of search results are provided in the Supplementary table 1). The structures 2XRC, 1Z8G, and 2OQ5 were the most closely related to the extracellular region of both proteins (Figure 1C). TMPRSS2 showed the best match with the 2XRC structure of Human complement factor I encoded by the CFI gene (Roversi et al., 2011), while TMPRSS4 with the 1Z8G structure belonging to another TTSP family member, hepsin (HPN), which is also known as TMPRSS1 (Herter et al., 2005). Hepsin and TMPRSS2 proteolytically cleave the Angiotensin-converting enzyme 2 (ACE2) in a similar manner (Heurich et al., 2014). The 2OQ5 structure is a part of the catalytic domain of TTSP family member DESC1 (Kyrieleis et al., 2007). DESC1 was shown to activate influenza viruses and coronaviruses in cell culture linked to host cell entry (Zmora et al., 2014). Most remaining HHPred hits belonged to structures of complement factors or blood coagulation and anticoagulation proteins (Table 1). It is noteworthy, that not a single structural match was obtained for the cytoplasmic region of both proteins, even when their amino acid sequences were queried alone.

**Table 1:**
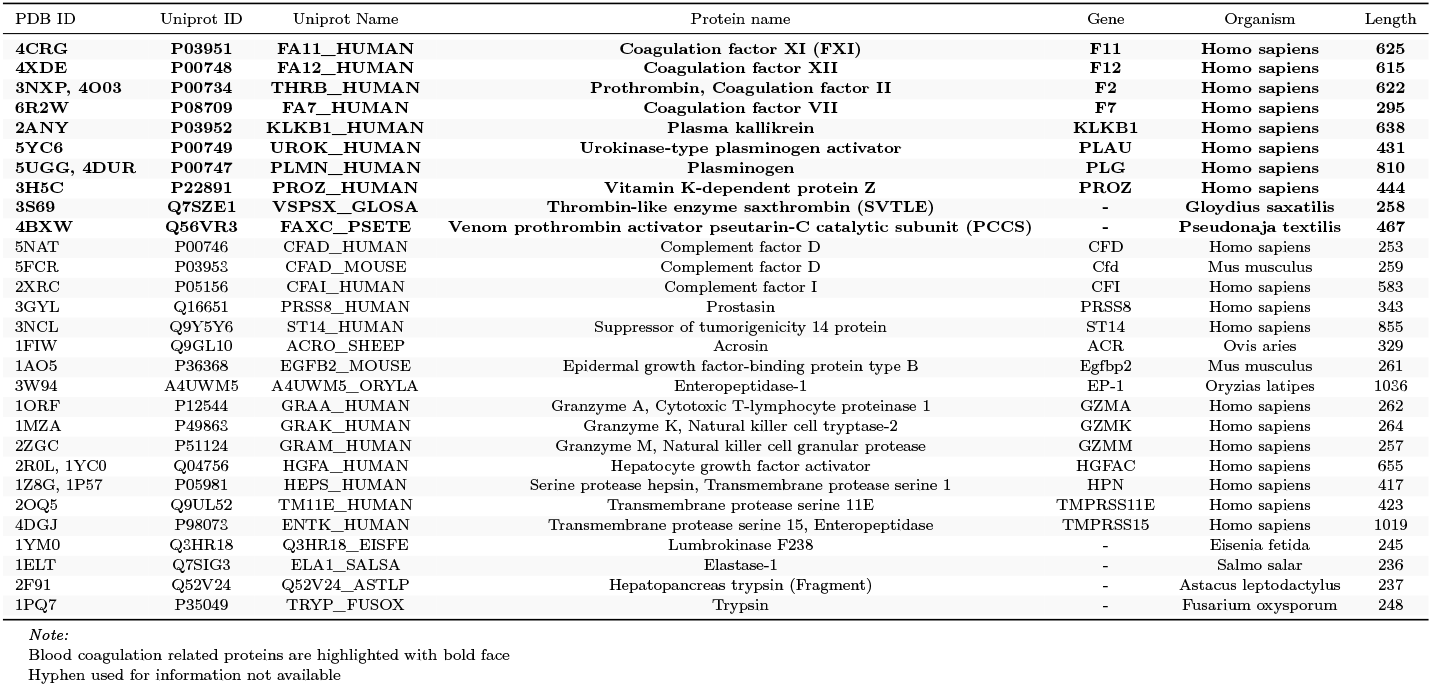
Structural homologs of TMPRSS2 and TPMRSS4 identified by HHPred and Phyre2

The high sequence similarity of TMPRSS2 and TMPRSS4 with known structures in the PDB database allowed modelling their 3D structure using Phyre2 web server (Kelley et al., 2015). In both sequences, about 86% of the residues were modelled at more than 90% confidence. The 67 and 61 residues at the N-terminal ends of TMPRSS2 and TMPRSS4, respectively, were modelled ab initio due to the lack of homology with known structures. Therefore, we removed the coordinates of the first 125 and 61 amino acids from the predicted structures of TMPRSS2 and TMPRSS4 due to their low confidence prediction. Figure 2 shows that SRCR and the serine-protease domains were modelled with high accuracy in both proteins with the preservation of clustered cysteines and the catalytic triad of serine-protease domain, respectively. A similar adjacent two-domain architecture is common in other TTSPs such as TMPRSS5 and Hepsin (Herter et al., 2005; Szabo and Bugge, 2008).

**Figure 2:**
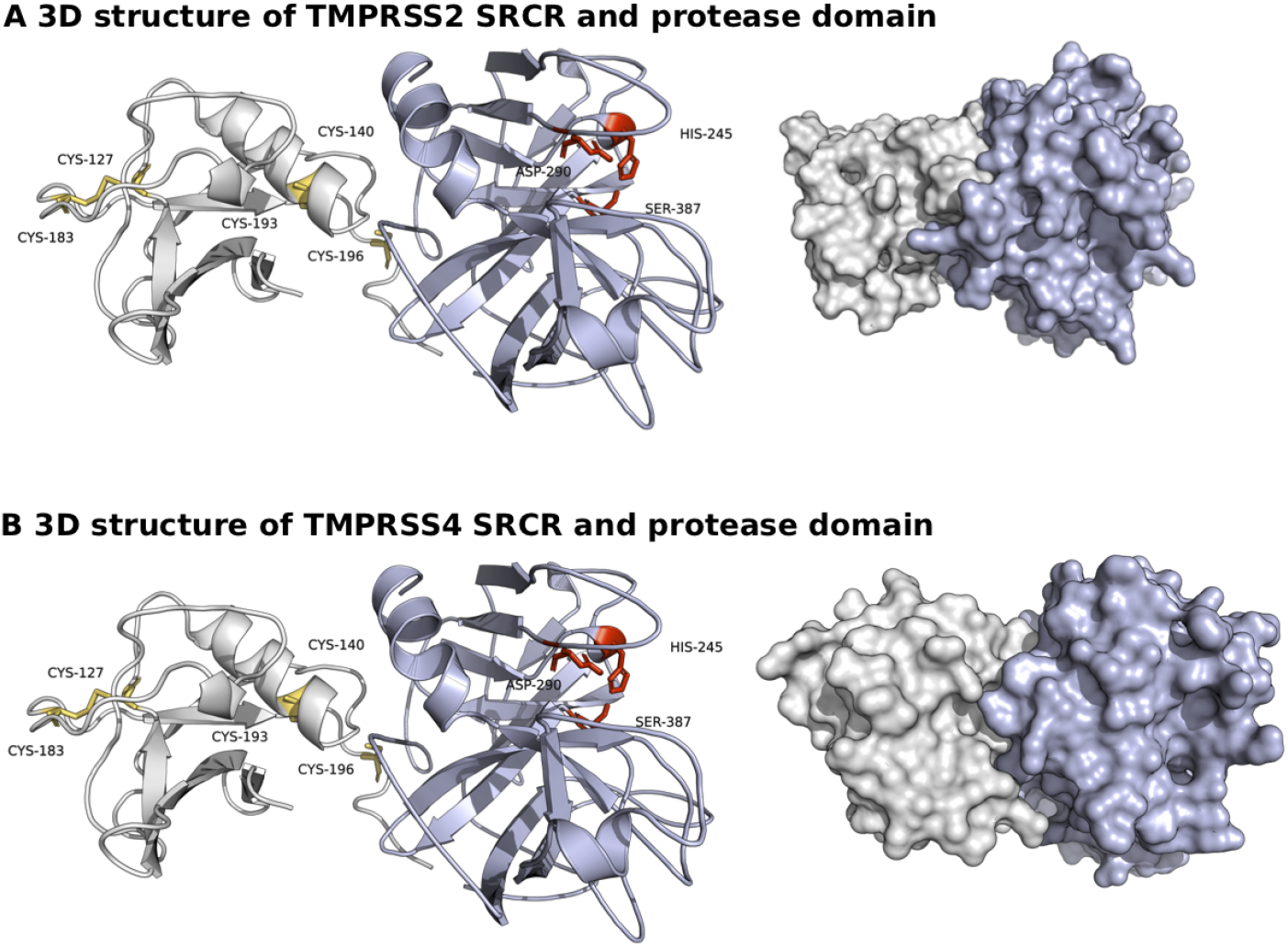
Predicted 3D structures of TMPRSS2 and TMPRSS4 SRCR and serine-protease domains. The figure shows structural decomposition of predicted proteins. Greek-key *β*-barrel fold and scavenger receptor cysteine-rich (SRCR) domains are shown in indigo and grey, respectively.

The template structures identified by Phyre2 (see Material and Methods) and the top 20 hits obtained by the HHPred search of TMPRSS2 and TMPRSS4 homologs correspond to a total of 36 structures of 29 proteins from 10 species. As expected, all of these correspond to known proteases and, strikingly, many of them are related to blood coagulation processes, both pro- and anticoagulation (10 of 29 proteins) (Table 1). These proteins include anticoagulation factor plasminogen, coagulation factor thrombin and plasma kallikrein. Moreover, five of the identified proteins were linked to immune functions, including the Complement factor D and I involved in an alternative immune response complement pathway, as well as the Granzymes GZMA, GZMK, and GZMM required for activation of caspase-independent cell death in cytotoxic T-cells and NK-cells (Ewen et al., 2012). We also observed that the six proteins encoded by the genes F11, F2, KLKB1, PLAU, ACR, and TMPRSS11E are the targets of SERPINA5, a plasma serine protease inhibitor with hemostatic roles as a procoagulant, anticoagulant and proinflammatory factor (Yang and Geiger, 2017).

We further analyzed the domain architecture of these proteins. The serine-protease domain is conserved in all homologs, and half of them are also accompanied by other domains, particularly those found in proteins related to blood coagulation (Figure 3). These findings prompted the possibility that TMPRSS2 and TMPRSS4 have functions in blood pro- and anticoagulation related processes. We believe that these functions could be performed by their serine protease domain alone as the thrombin-like snake venom serine protease (UniProt name, VSPSX_GLOSA) has only the protease domain and it shows strong blood coagulation activity *in vitro* (Figure 3) (Wei et al., 2007).

**Figure 3:**
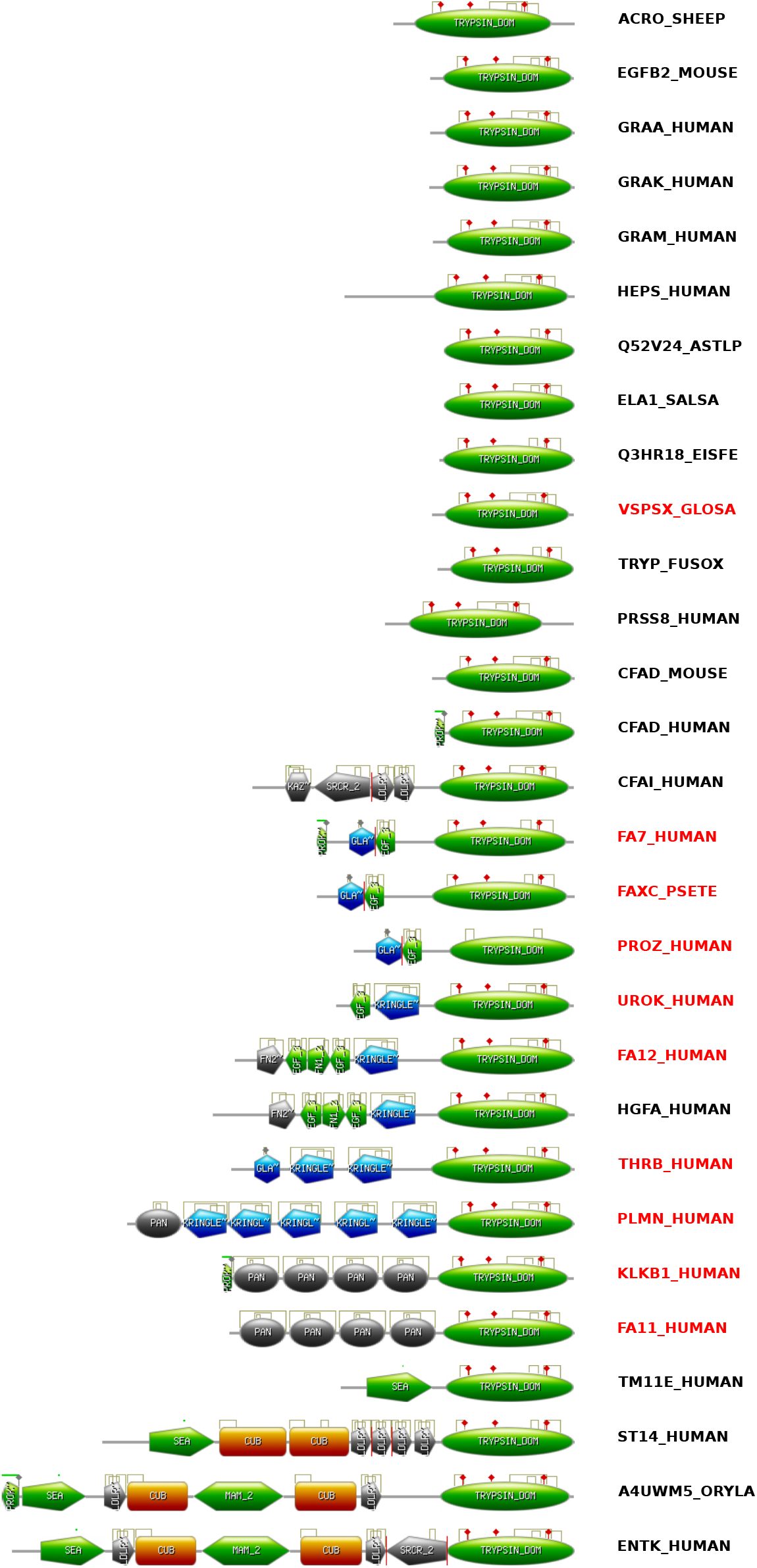
Domain architectures of TMPRSS2 and TMPRSS4 structural homologs. The figure shows domain architectures of 29 proteins from ProSite database. The active sites are shown with red diamonds and disulphide bridges with golden lines.

### 3.4. Structural homology of the predicted structure of TMPRSS2 and TMPRSS4 with plasminogen and enteropeptidases

We used the align module in PyMOL to compute the RMSD between backbone atoms of the predicted structures of TMPRSS2 and TMPRSS4 with each of the above-mentioned 36 structures. More than 30 structures showed high similarity with both structures with RMSD values of less than 1Å (Table 2). Among them, the PDB structure 5UGG containing a serine protease domain showed a striking superimposition with RMSD values of 0.563Å and 0.499Å with TMPRSS2 and TMPRSS4, respectively (Figure 4A, C). The 5UGG structure belongs to a plasminogen (Law et al., 2017). Further underscoring the significance of this finding is that along with the overall similarity of their chymotrypsin fold, the coordinates of the active site amino acid triad are almost identical between 5UGG and both predicted structures (Figure 4B, D). The second-best structural alignment of TMPRSS2 was with the 4DGJ structure with a RMSD value of 0.611Å, and TMPRSS4 with 3W94 with a RMSD value of 0.532Å (Table 2). The 4DGJ and 3W94 structures are representative of TTSP enteropeptidases such as the human TMPRSS15 which is found on the brush border membrane of epithelial cells in the duodenum and its homolog from the Japanese rice fish *Oryzias latipes* (UniProt accession A4UWM5), respectively. On the other hand, the TMPRSS4 SRCR domain shows structural similarity with the equivalent domain in hepsin (PDB 1Z8G, with a RMSD value of 0.169Å), while the TMPRSS2 SRCR domain was more similar to the chain D of the Complement factor I domain (PDB 2XRC with RMSD value of 0.458Å) (Table 2) (Herter et al., 2005; Roversi et al., 2011).

**Table 2:**
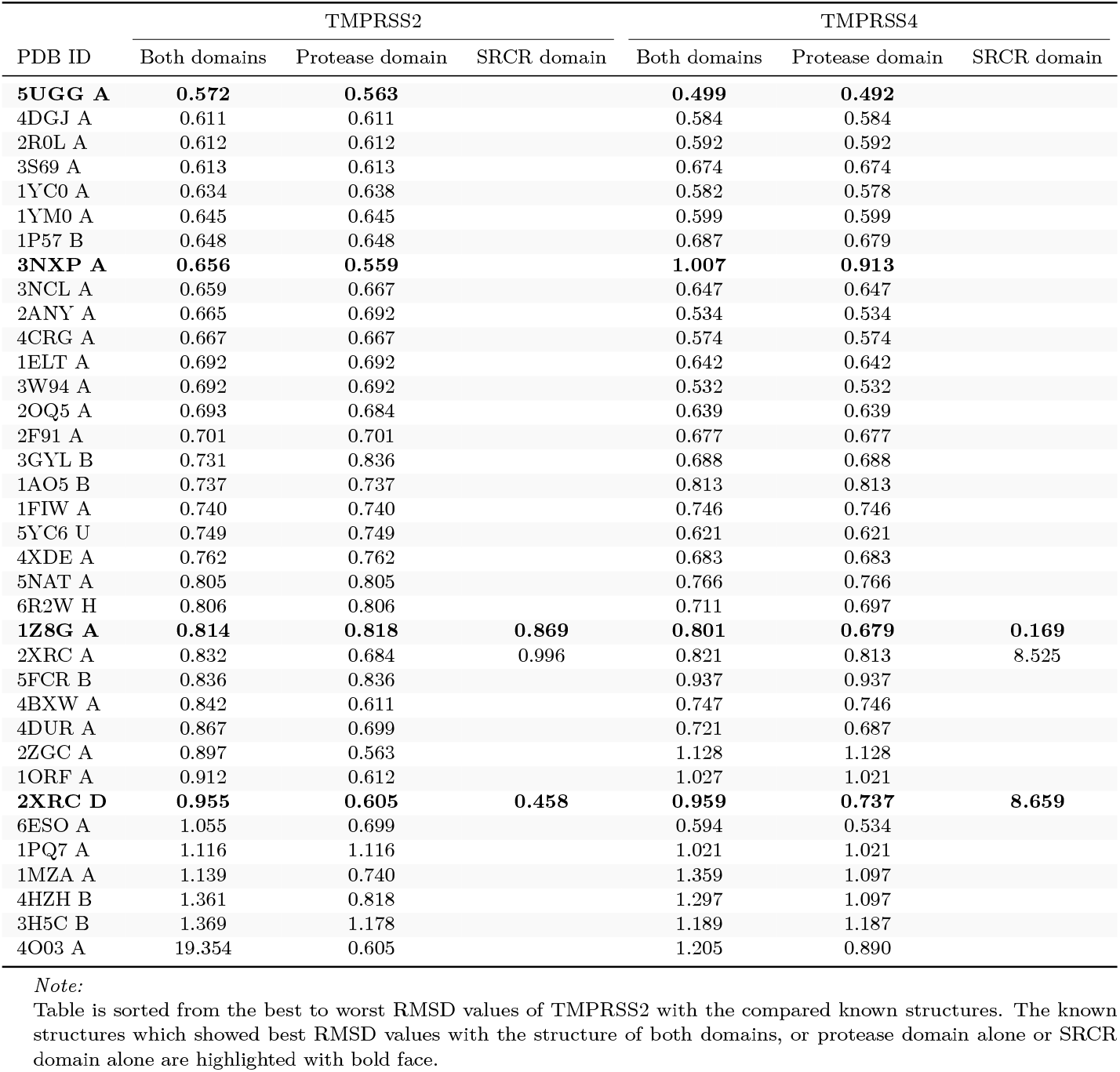
Root mean square deviations of predicted TMPRSS2 and TPMRSS4 3D structures with 36 known structural homologs

**Figure 4:**
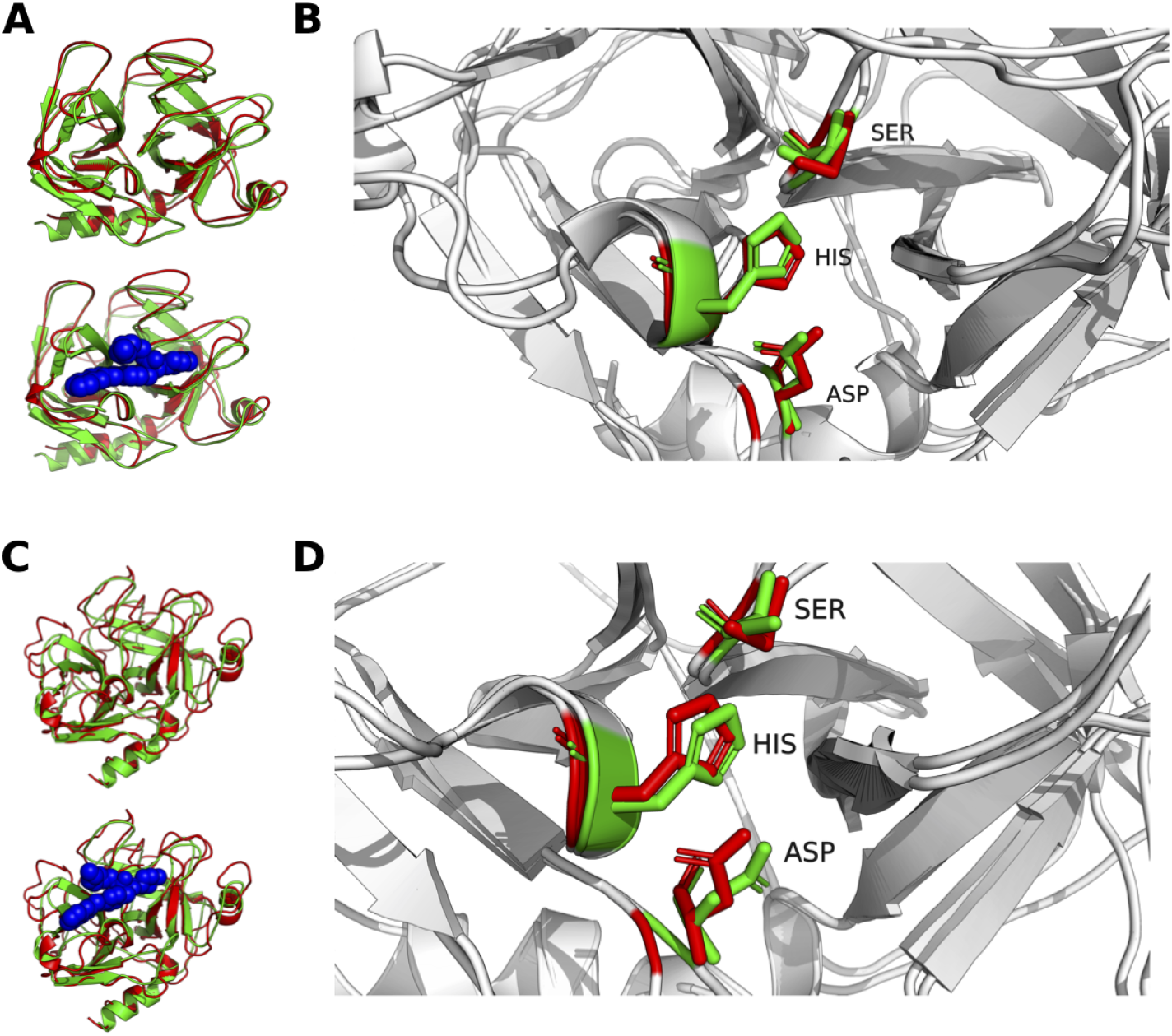
Superimposition of protease domains of predicted 3D structures of TMPRSS2 and TMPRSS4 with other known protease domains. Superimposition is shown of the protease domain of the 5UGG PDB structure of plasminogen with TMPRSS2 (A) and TMPRSS4 (B). TMPRSS2 and TMPRSS4 domains are shown in green while the 5UGG domain in red. Panel B and D shows the superimposition of the serine-protease catalytic triad (Ser-His-Asp) of TMPRSS2 and TMPRSS4 in green and 5UGG in red. The structure of 5UGG ligand (PDB ID 87M) is shown with blue spheres in (A) and (C).

Corroborating the above results with HHPred predictions suggest that TMPRSS2 and TMPRSS4 are likely to adopt a structural core similar to the two TTSP enteropeptidases, TMPRSS15 and Hepsin, the Complement factor I, and plasminogen since they were most closely related at the sequence and predicted structure level (Heissig et al., 2020; Lo et al., 2020; Risitano et al., 2020). These results are consistent with previous in-silico analyses of the predicted TMPRSS2 structure. Hepsin was used as a template to model TMPRSS2 structure in three previous studies (Chikhale et al., 2020; Idris et al., 2020; Rahman et al., 2020), while human TMPRSS15 and its Japanese rice fish ortholog structures were also used (Hempel et al., 2021; Huggins, 2020).

### 3.5. TMPRSS2 and TMPRSS4 may be inhibited by thrombin, plasma kallikrein and plasminogen inhibitors

The plasminogen serine protease domain structure showed the best overlap with TMPRSS2 and TMPRSS4 protease domains. Plasminogen dissolves the fibrin and dissolves blood clots (Storti and Szwast, 1982), and this activity has been shown to be inhibited by Aprotinin, a polypeptide consisting of 58 amino acid residues from bovine lung (Mahdy and Webster, 2004). Interestingly, Aprotinin also inhibits plasma kallikrein and thrombin, which are involved in blood coagulation and showed a reasonable structural match with TMPRSS2 and TMPRSS4 serine protease domains. Therefore, we assumed that their selective ligands can also inhibit the activity of TMPRSS2 and TMPRSS4. Structures of the complexes of 5UGG (plasminogen) and 4BXW (prothrombin activator) with their selective inhibitors 89M and OGJ, respectively are available in the PDB database (Law et al., 2017; Lechtenberg et al., 2013). In addition, we selected 22U and 75D molecules, which are selective inhibitors of thrombin (PDB, 2ZC9) and plasma kallikrein (PDB, 6O1S) (Baum et al., 2009; Partridge et al., 2019). The latter structures were treated as positive controls since they were not used as templates for TMPRSS2 and TMPRSS4 structure prediction by Phyre2. In addition, we selected the structure of 2ANY of the kallikrein protease family (Tang et al., 2005). These four inhibitor molecules were then used for docking with TMPRSS2, TMPRSS4, 4BXW, 5UGG, and 2ANY using the Glide docking program. We selected 4BXW, 5UGG, and 2ANY among the other structures since they represent thrombin, plasminogen, and kallikrein family serine protease structures and they had the lowest RMSD values with the TMPRSS2 and TMPRSS4 structures. The receptor grid was centered on the center of mass of the catalytic triad (Ser-His-Asp) of these proteins. Interestingly, most Glide docking scores for TMPRSS2 and TMPRSS4 with all four inhibitors were as good as with their original receptor molecules (Table 3). The predicted structures of TMPRSS2 and TMPRSS4 had binding energy values below −5 kcal/mol for all molecules except for TMPRSS4, which had a binding energy of around −8 kcal/mol for the inhibitors OGJ and 89M. Hence, TMPRSS2 scores reflected a close fit for all inhibitors suggesting that it has a 3D structure similar to blood coagulants as well as anticoagulant. However, TMPRSS4 showed a reasonably high selectivity only for the inhibitors of the blood coagulation factors thrombin and kallikrein. All these reported inhibitors are known to interfere with the catalytic triad of the serine proteases. As shown in Figure 5, although both TMPRSS2 and TMPRSS4 have the same number of bonds with both inhibitors, the latter imparts an extra charged interaction with OGJ using Lys287, which also forms a cation-pi and hydrogen bond with the 87M ligand, allowing a tighter binding.

**Table 3:** Glide docking scores (in kcal/mol) of blood coagulation related protein inhibitors with their known targets plasminogen, plasma kalikrein, prothrombin activator, and with predicted structures of TMPRSS2 and TMPRSS4.

| Protein name or PDB ID | Inhibitors |  |  |  |
| --- | --- | --- | --- | --- |
|  | OGJ<sup>a</sup> | 22U<sup>a</sup> | 89M<sup>b</sup> | 7SD<sup>c</sup> |
| TMPRSS2 | -4.851 | -5.123 | -4.332 | -4.602 |
| TMPRSS4 | -8.842 | -5.565 | -8.564 | -5.92 |
| 4BXW (Venom prothrombin activator pseutarin-C catalytic subunit) | -5.985 | -3.283 | -4.145 | -5.12 |
| 5UGG (Plasminogen) | -4.576 | -4.589 | -4.576 | -4.873 |
| 2ANY (Plasma kallikrein, light chain) | -4.45 | -5.11 | -4.238 | -5.489 |

*Note:*
TMPRSS2 and TMPRSS4 docking scores are based on predicted strcutures. 4BXW, 5UGG, and 2ANY are PDB structure identifiers, whose protease domains were used for docking. All these structures are part of the blood coagulation factors (annotated in brackets)
<sup>a</sup> Peptide like selective thrombin inhibitor;
<sup>b</sup> Plasminogen inhibitor;
<sup>c</sup> Plasma kalikrein inhibitor;

**Figure 5:**
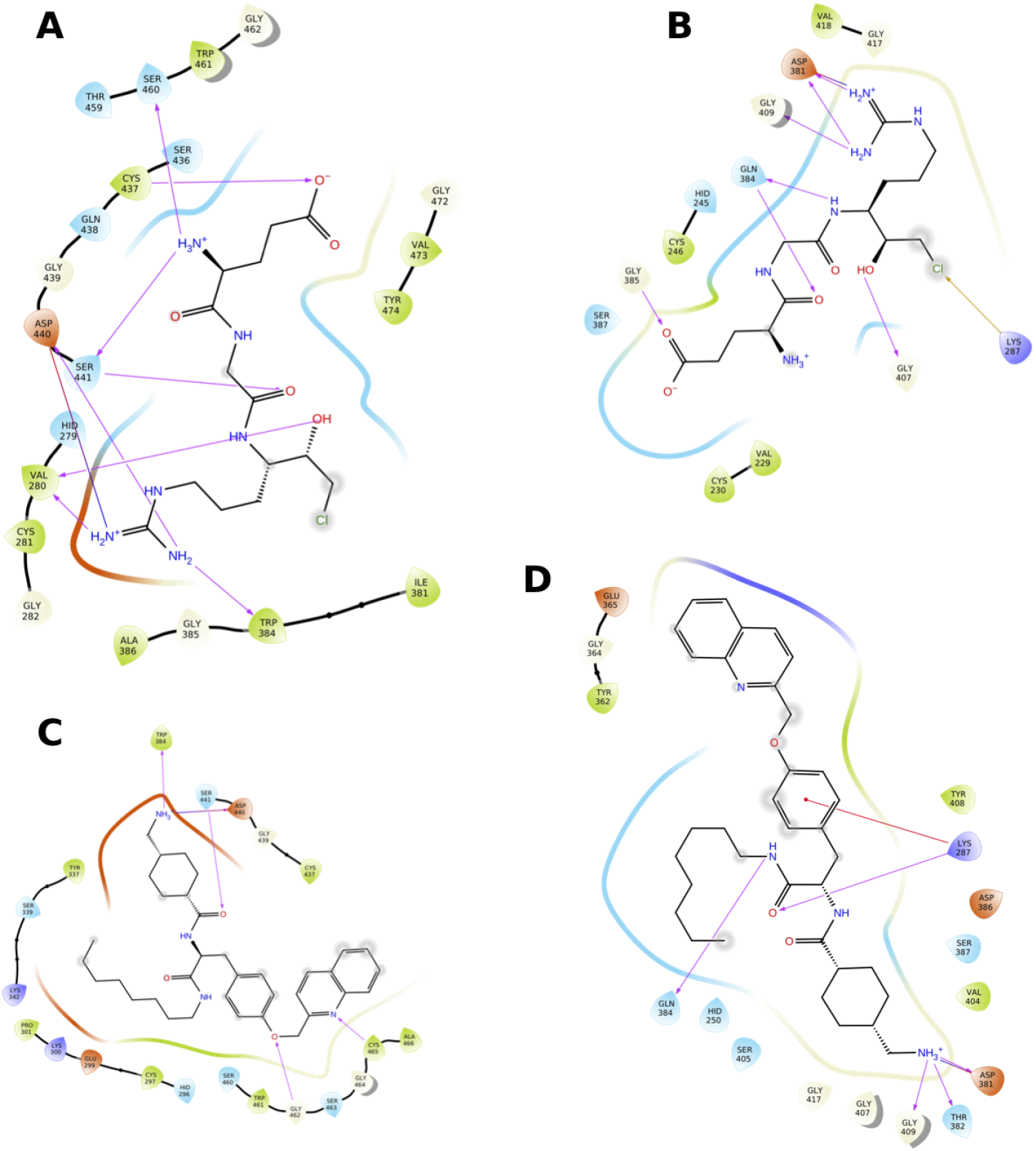
Docking of TMPRSS2 and TMPRSS4 with known protease inhibitors. Docking is shown of TMPRSS2 (A) and TMPRSS4 (B) with the peptide-like thrombin inhibitor OGJ and the plasminogen tranexamic acid-derived inhibitor 89M (C) and (D). The docking score of OGJ with TMPRSS2 was −4.851 (kcal/mol), and with TMPRSS4 was −8.980 (kcal/mol), whereas with the inhibitor 89M was −4.332 (kcal/mol) and −8.564 (kcal/mol), respectively. Hydrogen bonds are indicated by pink arrows and cation-pi interactions by red arrows.

In summary, these results suggest that the serine protease domains of the TMPRSS2 and TMPRSS4 are likely to have structural cores very similar to those of plasminogen, thrombin, and plasma kallikrein. Hence, they are likely to share effects as blood anticoagulant factors.

## 4. Discussion

The membrane-bound proteins TMPRSS2 and ACE2 are well known entry points of SARS-CoV-2. On the other hand, TMPRSS4 has not been well studied in the context of COVID19 pathogenesis. The TMPRSS4 expression on cell membranes have been shown to promote SARS-CoV-1 S protein driven cell-cell fusion similar to that of TMPRSS2 but without the cleavage of the S protein (Glowacka et al., 2011). This suggests that TMPRSS4 activates the S protein independently of its cleavage by an unknown molecular mechanism. TMPRSS2 and TMPRSS4 also activate hemagglutinin, which is indispensable for influenza virus infectivity in lungs (Chaipan et al., 2009). Moreover, both proteins have also been shown to assist infection of SARS-CoV-2 infection in the intestine (Zang et al., 2020). This evidence suggests that TMPRSS2 and TMPRSS4 perform related functions. Therefore, we analyzed TMPRSS2 and TMPRSS4 proteins at the sequence and their predicted structure levels and provide evidence supporting their related functions.

### 4.1. The N-terminal cytoplasmic region of TMPRSS2 is similar to the domain involved in protein trafficking

The N-terminal cytoplasmic region of 85 residues in TMPRSS2 is likely to adopt the structural fold (CATH ID: 1.10.185.10) found in Human T-cell Leukemia Virus Type II Matrix Protein (Dawson et al., 2017). This gag protein is a common feature of all retroviruses and is required for membrane localization of the assembling viral particle and subsequently remains associated to the inner surface of the membrane of the mature virion (Christensen et al., 1996). Interestingly, we also found a hit for another domain in this region belonging to the *β*-sandwich domain of the Sec23/24 superfamily (CATH ID: 2.60.40.1670). This domain is a prototype of sec23/24 proteins that are part of the multi-subunit complex COPII coat, which is responsible for the selective export of cargo proteins from the endoplasmic reticulum to the Golgi apparatus (Hughes and Stephens, 2008). Both domains seem to be involved in cargo trafficking. However, we did not find homologous sequences for this region in viral genome restricted searches using a PSI-BLAST at NCBI, neither using HHPred searches in the PDB database (Hildebrand et al., 2009; Johnson et al., 2008). This is likely due to the fact that structures are more conserved than the sequences and hence it is often observed that the same fold is adopted by proteins even though amino acid sequences differ substantially. Therefore, experimental analyses are required to address the functional aspects of both predicted domains considering their possible roles in viral assembly and trafficking (Christensen et al., 1996; Hughes and Stephens, 2008).

### 4.2. LDLRA domains of TMPRSS2 and TMPRSS4 may be invovled in viral entry via receptor-mediated endocytosis

The extracellular part of both proteins has highly conserved SCRC, LDLRA, and serine-protease domains in the same order. The LDLRA domains are unstructured and their binding of the calcium ion confers them structural integrity, which in turn allows their binding to cholesterols. The calcium binding site is conserved in both proteins suggesting that they can form an active LDLRA structure. Cholesterol is an essential component of the eukaryotic cell membranes in which it plays a critical role by maintaining their fluidity and hence the barrier between cell and environment. Membrane receptors having LDLRA domains bind LDLs that contain esterified cholesterol and carry them into cells after clustering in clathrin-coated pits by a receptor-mediated endocytosis (Daly et al., 1995). However, the function of the LDLRA domain in TMPRSS2 and TMPRSS4 is not clear as of now. It may normally bring LDLs inside cells through receptor-mediated endocytosis as it does with other proteins such as apolipoprotein E (Daly et al., 1995,Brown and Goldstein (1986)), which can be used for the host membrane remodelling. Assuming that viruses exploit host cell machinery to perform their tasks, TMPRSS2 mediated endocytosis can also import SARS-CoV-2 inside cells along with LDLs. Since endocytosis leaves no evidence of virus entry and it can avoid detection of its cargo by immunosurveillance, the endocytic pathway appears to be a common mechanism of host cell entry for many viruses. For example, herpes simplex virus 1 and human immunodeficiency virus 1 are capable of entering directly but often use the endocytic pathways for cell entry (Daecke et al., 2005; Miyauchi et al., 2009; Nicola et al., 2003). The membrane domains with high concentrations of cholesterols known as lipid rafts have also been shown to be targeted by viruses for cell entry (Lingwood and Simons, 2010; Simons and Ikonen, 1997). The importance of clathrin-endocytotic pathway, lipid rafts and the presence of ACE2 receptors in them has been confirmed in SARS-CoV-1 (Glende et al., 2008; Inoue et al., 2007; Wang et al., 2008) as well as in SARS-CoV-2 infections (Li et al., 2021; Nardacci et al., 2021). Furthermore, many RNA viruses harness endocytosis to traffic cholesterol from the remodeled membrane and extracellular medium to generate replication organelles, where cholesterol regulates viral polyprotein processing and genome replication (Ilnytska et al., 2013). The Cholesterol-25-hydroxylase converts cholesterol to 25-hydrocholesterol and depletes it from host membrane, which in turn has been shown to block SARS-CoV-2 membrane fusion thereby inhibiting infection in lung epithelial cells (Wang et al., 2020). In addition, higher levels of oxidized cholesterols could lead to the induction of a procoagulant state (Kim et al., 2020) or aggravate the formation of atherosclerotic plaques. This would in part explain the blood clotting and breathing problems observed in COVID19 patients (Gu et al., 2021). Therefore, TMPRSS2 and TMPRSS4 LDLRA domains must be studied further.

### 4.3. Serine-protease domains of TMPRSS2 and TMPRSS4 show significant similarity with plasminogen and blood coagulation factors

The TMPRSS2 and TMPRSS4 3D structures are not available. Hence, we first identified their structural homologs using HHPred, and also modelled 3D structures using Phyre2. We identified 29 structural homologs from 10 species with both approaches. These 3D structures were mapped to the SCRC and serine-protease in both proteins, but no match was found for the LDLRA domain. When we analyzed further the domain architectures of the homologs, we found that the serine-protease domain is conserved in all, and half of them are also accompanied by other domains, particularly those found in proteins related to blood coagulation and anticoagulation. As discussed earlier, the TMPRSS2 and TMPRSS4 serine protease domains are present at their C-termini. The TMPRSS2 domain, which is tethered to the outer face of the cell membrane, activates the S protein of SARS-CoV-2 for host cell entry. Among the serine-protease domains of 29 proteins, that of plasminogen showed a striking superimposition with TMPRSS2 and TMPRSS4 and the coordinates of their active site amino acid triad were almost identical. It is also shown that the plasminogen shows 95% identity within the S1–S1’ subsites of TMPRSS2, which are used for cleavage of the SARS-Cov-2 spike protein, and 64.71% within S4–S4’ subsites, which were the highest among 14 serine proteases including TMPRSS15 which was selected for homology modelling of the TMPRSS2 structure (Huggins, 2020). This suggests that the serine-protease domains of the TMPRSS2 and TMPRSS4 are likely to have a protease activity similar to that of plasminogen, which dissolves the fibrin of blood clots (Storti and Szwast, 1982). Hence, these findings suggest that TMPRSS2 and TMPRSS4 serine protease domains have similar catalytic properties to those of blood clotting factors further supporting the notion that they are likely involved in blood coagulation and anticoagulation through common mechanisms involving plasminogen, plasma kallikrein and thrombin. Similar to these three blood coagulation related factors, the protease domains of TMPRSS2 and TMPRSS4 are also extracellular and they are made also snthesized in an inactive zymogen form. Interestingly, TMPRSS4 directly activates the urokinase-type plasminogen activator (pro-uPA) encoded by the PLAU gene through its proteolytic activity, which in turn can cleave zymogen plasminogen to form the active enzyme plasmin (Min et al., 2014). This suggests a potential role of TMPRSS4 in blood clot resolution upstream of plasminogen and pro-uPA, which is one of the TMPRSS2 and TMPRSS4 structural homologs identified in this study. It is noteworthy that pro-uPA and plasminogen are both ligands for the LDLRA domain containing protein families (Liu et al., 2001). Hence additional studies are warranted to determine whether TMPRSS4 activates pro-uPA which in turn converts plasminogen to plasmin to resolve blood clots. These sequential reactions are likely to be dependent on the LDLRA domain of TMPRSS4 and we believe TMPRSS2 also performs similar functions.

### 4.4. Plasminogen, thrombin, and plasma kallikrein inhibitors show selective binding to TMPRSS2 and TMPRSS4 serine-protease active sites

Aprotinin has been shown to inhibit plasminogen, thrombin, and plasma kallikrein. Therefore, we assumed that the inhibitors of these proteins can also inhibit TMPRSS2 and TMPRSS4. To test this, we used 3D structures of four selective inhibitors and performed docking analysis with the protease active sites of TMPRSS2, TMPRSS4, prothrombin activator, plasminogen, and plasma kallikrein structures. As expected, the Glide docking scores with all four inhibitors for TMPRSS2, and to a lesser extent for TMPRSS4, were as good as with their original molecules suggesting that these inhibitors are likely to impair TMPRSS2 and TMPRSS4 protease activity. Moreover, Aprotinin, a polypeptide consisting of 58 amino acid residues from bovine lung inhibits plasminogen, plasma kallikrein and thrombin (Mahdy and Webster, 2004). There is also evidence that it specifically inhibits other serine-proteases including TMPRSS2 and TMPRSS4 that cleave hemagglutinin protein of Influenza virus, and aerosol inhalation of Aprotinin is used for the treatment of patients with mild-to-moderate influenza infections (Ovcharenko and Zhirnov, 1994; Zhirnov et al., 2011). Aprotinin has been suggested to inhibit TMPRSS2 as well (Shen et al., 2017).

### 4.5. TMPRSS2 and TMPRSS4 may function at high-temperature during immune response owing to their serine-protease domain

Strikingly, the 29 structural homologs of TMPRSS2 and TMPRSS4 we detected can be grouped into a few related functions including 10 in blood coagulation related processes, 5 in immune functions, and the rest in alternative immune response complement pathway, and activation of caspase-independent cell death in cytotoxic T-cells and NK-cells (Ewen et al., 2012). Interestingly, the five proteins encoded by the genes F11, F2, KLKB1, PLAU, and ACR, and also TMPRSS member TMPRSS11E are the targets of SERPINA5, a plasma serine protease inhibitor with hemostatic roles as a procoagulant, anticoagulant and proinflammatory factor (Yang and Geiger, 2017). It is highly possible that TMPRSS2 and TMPRSS4 genes are also targets of SERPINA5. Additionally, it is known that dexamethasone induces expression of the beta isoform of GZMA and represses expression of its alpha isoform, upon binding of the glucocorticoid receptor (Ruike et al., 2007), ultimately leading to apoptotic cell death (Myoumoto et al., 2007). Interestingly, septic shock is an inflammatory response which causes excessive cell death, and critical COVID19 patients have been shown to recover using Dexamethasone (RECOVERY Collaborative Group et al., 2021). Furthermore, anticoagulants have been used to treat COVID19 patients with good success (Levi et al., 2020; Violi et al., 2020).

One of the hallmarks of immune response is fever, especially upon infections by viruses or other pathogens. Presumably TMPRSS2 and TMPRSS4 along with their 29 homologs, necessitate to remain stable and functional at high temperature during fever. Intriguingly, their serine-protease domain is a prototype feature of the high temperature requirement A (HtrA) protein family, whose members act as chaperones and are responsible for maintaining protein quality at high temperature (Clausen et al., 2002). The HtrA family is present in the three domains of life and hence, the HtrA mediated cellular response to high temperature is universally conserved (Muley et al., 2019). Null mutants of *Escherichia coli* HtrA do not survive at elevated temperature due to instability of temperature sensitive proteins due to the lack of protein quality control measures mediated by HtrA (Lipinska et al., 1990). These observations indicate that the serine-protease domains are likely selected as a part of blood coagulation and anticoagulation, immune response, and alternative complement pathways to perform their functions efficiently during immune responses and keep normal hemostasis at high temperatures. Hence, the allele frequency and nucleotide sequences of these protein coding genes can be expected to vary among the populations originally adapted to cold, hot, and temperate zones (Wang et al., 2020). This could be one of the important factors involved in diverse rates of dispersion of SARS-CoV-2 infection in countries with different populations and temperatures. One cannot ignore the diverse genetic makeup of the world’s population influenced by the surrounding environment, and TMPRSS2 and TM-PRSS4 should be further studied in this regard especially their serine-protease domain.

## 5. Conclusions

Our in-silico analysis based on the predicted structures of TMPRSS2 and TMPRSS4 allowed the identification of structural homologs many of which are involved in blood coagulation, immune response, and proteolysis, which are important in the context of immune functions. The similarity of the proteolytic domains of TMPRSS2 and TMPRSS4 to that of the blood clotting factors suggest that their catalytic properties are similar as well. Indeed, the tight docking of known inhibitors for these factors to the catalytic sites of TMPRSS2 and TMPRSS4 strongly suggest that their activity is also inhibited. This would in part explain why anticoagulant treatments that have been used to treat COVID19 patients have had good success. In addition to treating the clotting problems in these patients, it is expected that the inhibition of the extracellular domains of TMPRSS2 and TMPRSS4 would inhibit their proteolytic effect on ACE2 thus limiting virus entry into the target cells. Moreover, inhibition of these proteins in platelets might also limit the thrombotic effects of SARS CoV-2 (Zhang et al., 2020). Hence, our studies shed light on a novel mechanism by which anticoagulant treatments might act to limit the severity of COVID19 inking LDL and clotting factors and offer an avenue for further exploration of therapeutic approaches for this disease that is affecting the whole world population.

---

**Supplementary table 1:** Structural homologs of TMPRSS2 and TPMRSS4 sequences identified using HHPred

| PDB ID | Prob<sup>a</sup> | E-value<sup>b</sup> | Score<sup>c</sup> | SS<sup>d</sup> | Query HMM<sup>e</sup> | Template HMM<sup>f</sup> |
| --- | --- | --- | --- | --- | --- | --- |
| **TMPRSS2 sequence search hits** |  |  |  |  |  |  |
| 2XRC A | 100.0 | 5.5E-30 | 268.4 | 345 | 110-491 | 201-558(565) |
| 1Z8G A | 100.0 | 1.9E-27 | 235.6 | 349 | 140-492 | 1-363(372) |
| 2OQ5 A | 99.9 | 6.3E-25 | 200.2 | 230 | 256-489 | 1-231(232) |
| 4DGJ A | 99.9 | 9.9E-25 | 199.1 | 233 | 256-489 | 1-235(235) |
| 4BXW A | 99.9 | 2.5E-24 | 216.9 | 344 | 117-491 | 7-412(423) |
| 6R2W H | 99.9 | 3.4E-24 | 197.6 | 234 | 256-491 | 1-237(249) |
| 1P57 B | 99.9 | 3.8E-24 | 198.0 | 236 | 256-492 | 1-246(255) |
| 3W94 A | 99.9 | 4.1E-24 | 194.5 | 233 | 256-489 | 1-235(235) |
| 2ANY A | 99.9 | 9.5E-24 | 192.9 | 236 | 256-492 | 1-239(241) |
| 3H5C B | 99.9 | 1.1E-23 | 203.6 | 296 | 119-489 | 1-317(317) |
| 4XDE A | 99.9 | 6.8E-24 | 197.5 | 237 | 255-492 | 2-247(257) |
| 4CRG A | 99.9 | 7.6E-24 | 192.9 | 234 | 256-490 | 1-237(238) |
| 3NCL A | 99.9 | 9.1E-24 | 193.3 | 232 | 256-489 | 1-240(241) |
| 5NAT A | 99.9 | 1.3E-23 | 192.2 | 230 | 256-492 | 1-231(232) |
| 5FCR B | 99.9 | 1.8E-23 | 191.0 | 231 | 256-492 | 1-232(234) |
| 3GYL B | 99.9 | 1.3E-23 | 195.3 | 235 | 256-490 | 1-243(261) |
| 2F91 A | 99.9 | 1.7E-23 | 191.5 | 231 | 256-489 | 1-237(237) |
| 1ELT A | 99.9 | 1.6E-23 | 192.0 | 228 | 256-488 | 1-236(236) |
| 5YC6 U | 99.9 | 1.7E-23 | 192.1 | 234 | 256-489 | 1-246(246) |
| 2R0L A | 99.9 | 2E-23 | 192.7 | 236 | 256-492 | 1-242(248) |
| **TMPRSS4 sequence search hits** |  |  |  |  |  |  |
| 1Z8G A | 100.0 | 1.1E-33 | 270.1 | 340 | 96-436 | 1-362(372) |
| 2XRC A | 100.0 | 1.2E-29 | 256.1 | 372 | 60-436 | 26-558(565) |
| 2OQ5 A | 100.0 | 5.3E-26 | 200.3 | 229 | 205-434 | 1-231(232) |
| 2R0L A | 99.9 | 1.7E-25 | 199.5 | 229 | 205-436 | 1-241(248) |
| 1ORF A | 99.9 | 2.4E-25 | 197.1 | 229 | 205-437 | 1-234(234) |
| 4XDE A | 99.9 | 4.8E-25 | 198.2 | 233 | 203-436 | 1-246(257) |
| 1P57 B | 99.9 | 1.9E-24 | 193.2 | 231 | 205-436 | 1-245(255) |
| 5NAT A | 99.9 | 2.3E-24 | 190.5 | 225 | 205-435 | 1-229(232) |
| 1YC0 A | 99.9 | 2.6E-24 | 196.4 | 242 | 195-437 | 21-277(283) |
| 5YC6 U | 99.9 | 1.7E-24 | 192.0 | 230 | 205-434 | 1-246(246) |
| 1AO5 B | 99.9 | 6.3E-25 | 194.4 | 224 | 205-436 | 1-236(237) |
| 5UGG A | 99.9 | 6.6E-24 | 189.4 | 239 | 194-436 | 6-251(251) |
| 2ZGC A | 99.9 | 3.7E-24 | 190.2 | 229 | 205-436 | 1-231(240) |
| 1YM0 A | 99.9 | 2.6E-24 | 190.9 | 228 | 205-436 | 1-238(238) |
| 1FIW A | 99.9 | 3.4E-24 | 196.3 | 232 | 205-436 | 1-251(290) |
| 4DGJ A | 99.9 | 3.2E-24 | 189.1 | 228 | 205-434 | 1-235(235) |
| 1MZA A | 99.9 | 4.1E-24 | 189.4 | 228 | 204-435 | 2-236(240) |
| 4BXW A | 99.9 | 4.6E-24 | 207.3 | 241 | 194-437 | 159-413(423) |
| 3S69 A | 99.9 | 2.3E-24 | 190.8 | 226 | 205-436 | 1-227(234) |
| 1PQ7 A | 99.9 | 3.8E-24 | 187.3 | 224 | 205-433 | 1-224(224) |

*Note:*
TMPRSS2 and TMPRSS4 protein sequences were searched against the entire PDB database using HHPred web-server. The top 20 hits for each protein are reported in the table with their PDB and chain identifiers.
<sup>a</sup> Probability of target to be a true positive;
<sup>b</sup> The number of hits one can expect by chance with a score better than the one for the target when scanning the database;
<sup>c</sup> Raw sequence similarity score;
<sup>d</sup> Secondary structure similarity score between query and target;
<sup>e</sup> Range of aligned match states from query HMM;
<sup>f</sup> Range of aligned match states from target HMM;

**Supplementary figure 1:**
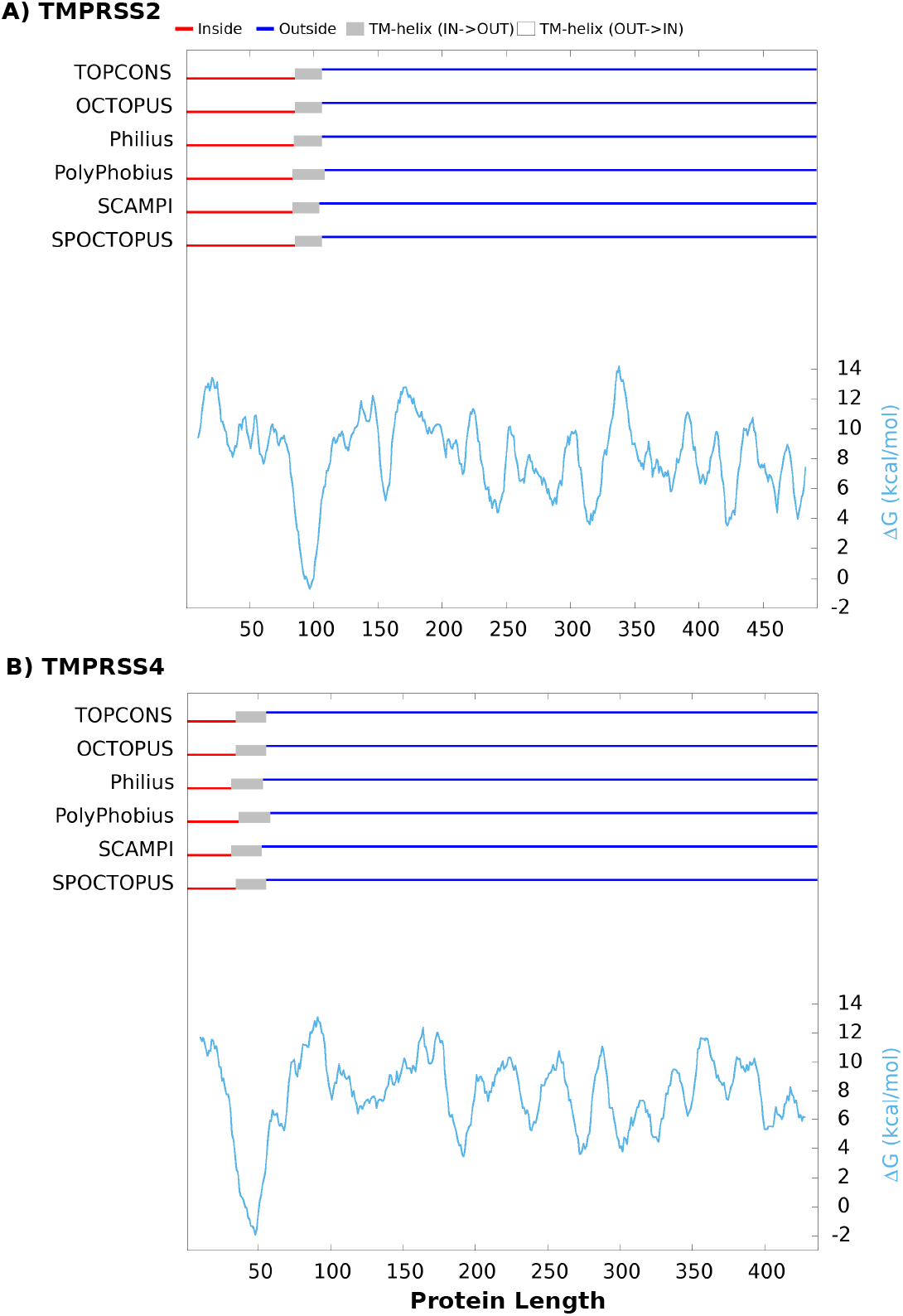
The position of membrane helix in TMPRSS2 and TMPRSS4. The combined analysis report obtained from the TOPCONS web-server is shown, in which lower G (free energy) values represent amino acids that are likely to be part of the trans-membrane helix. The thick red and blue lines represent the inside and outside topology of the protein, respectively. A transmembrane helix is predicted at the N-termini in TMPRSS2 (A) and TMPRSS4 (B) protein sequences.

**Supplementary figure 2:**
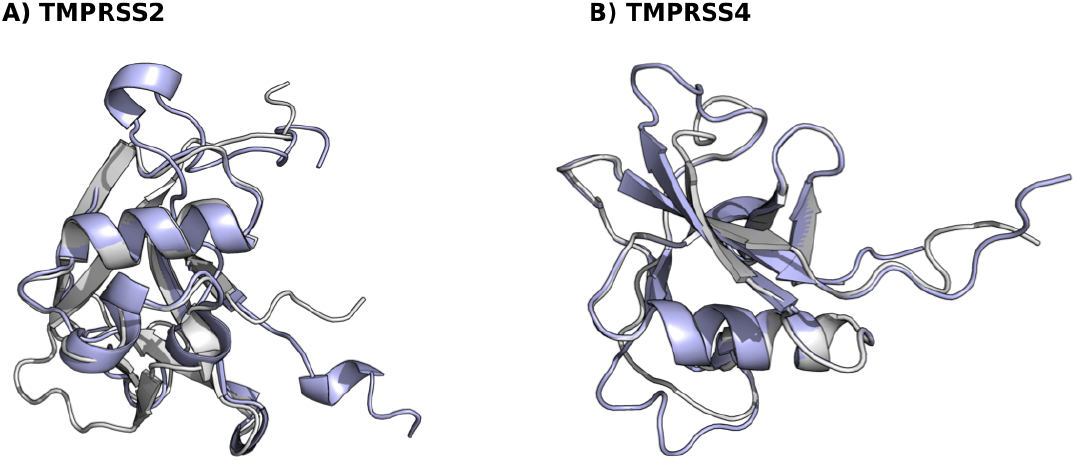
Superimposition of the protease domain of predicted 3D structures of TMPRSS2 and TMPRSS4 with the scavenger receptor cysteine-rich (SRCR) domain of 1Z8G PDB structure. TMPRSS2 (A) and TMPRSS4 (B) SRCR domains are shown in purple while 1Z8G is shown in grey.

